# InvL, an invasin-like adhesin, is a type II secretion system substrate required for *Acinetobacter baumannii* uropathogenesis

**DOI:** 10.1101/2022.02.01.478765

**Authors:** Clay D. Jackson-Litteken, Gisela Di Venanzio, Nguyen-Hung Le, Nichollas E. Scott, Bardya Djahanschiri, Jesus S. Distel, Evan J. Pardue, Ingo Ebersberger, Mario F. Feldman

**Affiliations:** Department of Molecular Microbiology, Washington University School of Medicine, St. Louis, Missouri 63110, United States; Department of Microbiology and Immunology, University of Melbourne at the Peter Doherty Institute for Infection and Immunity, Parkville, VIC 3010, Australia; Applied Bioinformatics Group, Inst. of Cell Biology and Neuroscience, Goethe University Frankfurt, Frankfurt am Main 60438, Germany; Senckenberg Biodiversity and Climate Research Centre (S-BIKF), Frankfurt am Main 60325, Germany; LOEWE Center for Translational Biodiversity Genomics (TBG), Frankfurt am Main 60325, Germany

## Abstract

*Acinetobacter baumannii* is an opportunistic pathogen of growing concern, as isolates are commonly multidrug resistant. While *A. baumannii* is most frequently associated with pulmonary infections, a significant proportion of clinical isolates come from urinary sources, highlighting its uropathogenic potential. The <u>t</u>ype <u>II</u> <u>s</u>ecretion <u>s</u>ystem (T2SS) of commonly used model *Acinetobacter* strains is important for virulence in various animal models, but the potential role of the T2SS in <u>u</u>rinary <u>t</u>ract <u>i</u>nfection (UTI) remains unknown. Herein, we used a <u>c</u>atheter-<u>a</u>ssociated UTI (CAUTI) model to demonstrate that a modern urinary isolate, UPAB1, requires the T2SS for full virulence. A proteomic screen to identify putative UPAB1 T2SS effectors revealed an uncharacterized lipoprotein with structural similarity to the intimin-invasin family, which serve as <u>t</u>ype <u>V</u> <u>s</u>ecretion <u>s</u>ystem (T5SS) adhesins required for the pathogenesis of several bacteria. This protein, designated InvL, lacked the β-barrel domain associated with T5SSs, but was confirmed to require the T2SS for both surface localization and secretion. This makes InvL the first identified T2SS effector belonging to the intimin-invasin family. InvL was confirmed to be an adhesin, as the protein bound to extracellular matrix components and mediated adhesion to urinary tract cell lines *in vitro*. Additionally, the *invL* mutant was attenuated in the CAUTI model, indicating a role in *Acinetobacter* uropathogenesis. Finally, bioinformatic analyses revealed that InvL is present in nearly all clinical isolates belonging to international clone 2, a lineage of significant clinical importance. In all, we conclude that the T2SS substrate InvL is an adhesin required for *A. baumannii* uropathogenesis.

**IMPORTANCE:** While pathogenic *Acinetobacter* can cause various infections, we recently found that 20% of clinical isolates come from urinary sources. Despite the clinical relevance of *Acinetobacter* as a uropathogen, few virulence factors involved in urinary tract colonization have been defined. Herein, we identify a novel type II secretion system effector, InvL, which is required for full uropathogenesis by a modern urinary isolate. Though InvL has predicted structural similarity to the intimin-invasin family of autotransporter adhesins, InvL is predicted to be anchored to the membrane as a lipoprotein. Similar to other invasin homologs however, we demonstrate that InvL is a bona fide adhesin capable of binding extracellular matrix components and mediating adhesion to urinary tract cell lines. In all, this work establishes InvL as an adhesin important for *Acinetobacter*’s urinary tract virulence, and represents the first report of a type II secretion system effector belonging to the intimin-invasin family.

## INTRODUCTION

*Acinetobacter* spp. belonging to the *<u>A</u>cinetobacter <u>c</u>alcoaceticus–Acinetobacter <u>b</u>aumannii* (ACB) complex are opportunistic pathogens that cause diverse infections, with *A. <u>b</u>aumannii* representing the most common species associated with human infection (1–3). While *A. baumannii* infections can be both community and hospital acquired, nosocomial infections in critically ill patients are common with mortality rates reported as high as 84.3% in individuals requiring mechanical ventilation (4–7). These high mortality rates can be largely attributed to increasing multidrug resistance among *A. baumannii* isolates (6, 8). In fact, *A. baumannii* exhibits the highest prevalence of multidrug resistance among Gram-negative pathogens, leading the World Health Organization to classify the bacterium as a highest priority pathogen for research and development of new treatments (8, 9). The respiratory tract is the most common site of infection by *A. baumannii*, with approximately 40% of the isolates derived from pulmonary infections (2, 3, 10, 11). It should be noted though that 20% of *A. baumannii* isolates come from <u>u</u>rinary <u>t</u>ract <u>i</u>nfections (UTIs), highlighting the bacterium’s uropathogenic capacity (10, 11). However, despite the public health relevance of *A. baumannii* as a uropathogen, few virulence factors required specifically for UTI have been defined (10).

Bacterial virulence relies on the interaction of bacterial-derived proteins with the infected host. Gram-negative bacteria have evolved to encode complex secretion systems to transport proteins across the bacterial envelope (12). The <u>t</u>ype <u>II</u> <u>s</u>ecretion <u>s</u>ystem (T2SS) is widely distributed in Gram-negative bacteria and has been implicated in the virulence of several pathogens (13–15). T2SS effectors are translocated from the cytoplasm to the outer membrane or into the extracellular milieu in two steps. First, the effector is transported across the inner membrane into the periplasm via the general <u>sec</u>retory (Sec) pathway or the <u>t</u>win <u>a</u>rginine <u>t</u>ranslocation (TAT) pathway (16, 17). Second, the effector is extruded through the outer membrane secretin (designated GspD herein) via the assembly of a pseudopilus (18–22). In *Acinetobacter*, the T2SS is responsible for secretion of several proteins, including the lipases, LipA, LipH, and LipAN, the protease, CpaA, and the <u>γ</u>-<u>g</u>lutamyltransferase, GGT (23–28). For A. *baumannii* strain ATCC 17978, a meningitis isolate from 1951, mutation of the T2SS results in attenuation in bacteremia and pneumonia models of infection (23, 29). Additionally, mutation of the T2SS in *Acinetobacter nosocomialis* strain M2 and A. *baumannii* strain AB5075, musculoskeletal isolates from 1996 and 2008, respectively, results in decreased bacterial burden in a pneumonia model of infection (24, 28). Furthermore, the *Acinetobacter* effectors LipAN of strain ATCC 17978 and CpaA of strain M2 are required for full virulence in a pneumonia model, and GGT of strain AB5075 is required for full virulence in a bacteremia model (23, 27, 28). While the T2SS and associated effectors clearly have roles in pathogenesis in multiple models of infection, the potential function of the T2SS in *Acinetobacter* uropathogenesis has not been investigated. Additionally, T2SS-dependent secretomes can differ significantly between strains, and these dissimilarities can lead to strain-dependent differences in virulence potential mediated by the T2SS (24, 28, 29). Therefore, evaluation of the T2SS in diverse clinical isolates could reveal previously unrecognized effectors which may serve as novel *Acinetobacter* virulence factors.

Intimate interaction of bacteria with host tissues at the site of infection is often required for pathogenesis. Indeed, adherence to and/or invasion of host epithelial cells, mediated by bacterial adhesins, is an essential process for the virulence of several pathogens (30). Many adhesins are displayed on the bacterial surface in a <u>T</u>ype <u>V</u> <u>s</u>ecretion <u>s</u>ystem (T5SS)-dependent manner (31). These T5SSs, also termed autotransporters (ATs), are proteins consisting of two general components, a β-barrel domain, which is inserted into the outer membrane, and a passenger domain, which uses the β-barrel for transportation to the outer surface of the bacteria (32–34). The intimin-invasin family of ATs specifically are adhesins that consist of an N-terminal β-barrel domain and a C-terminal passenger domain containing multiple immunoglobulin (IG)-like domains often capped by a lectin-like domain (35, 36). Intimin is encoded by many pathogens such as *Escherichia coli* and functions by binding to a <u>t</u>ype <u>III</u> <u>s</u>ecretion <u>s</u>ystem (T3SS) effector, <u>t</u>ranslocated intimin receptor (TIR), which is inserted into the host membrane (37, 38). Invasin (InvA), encoded by *Yersinia enterocolitica* and *Yersinia pseudotuberculosis*, functions by binding directly to host β1-integrins, leading to a host cytoskeletal rearrangement and subsequent internalization of the bacteria (39, 40). Importantly, deletion of genes encoding intimin-invasin family proteins results in attenuation in animal models of infection (35, 36). Despite this key role that intimin-invasin family proteins play in the pathogenesis of other bacteria, potential homologs have not been identified in *Acinetobacter*.

In this work, we aimed to interrogate the function of the T2SS in the recent *A. baumannii* urinary clinical isolate, UPAB1, using a murine <u>c</u>atheter-<u>a</u>ssociated <u>UTI</u> (CAUTI) model and define effectors potentially required for virulence (10). In this pursuit, we identify a novel T2SS effector/intimin-invasin family protein important for *Acinetobacter* uropathogenesis.

## RESULTS

### The UPAB1 T2SS mutant is attenuated in the CAUTI model

To test the role of the T2SS in uropathogenesis, we generated a UPAB1 mutant strain lacking *gspD*, the gene encoding for the outer membrane secretin that is essential for T2SS function. This strain was subsequently employed in the murine CAUTI model as previously described (10). Briefly, a small piece of silicon tubing (catheter) was inserted into the urethra, and mice were transurethrally inoculated with WT, Δ*gspD*, or complemented (*gspD+*) strains. 24 h postinfection, mice were sacrificed, and bacteria adhered to the catheters (**Fig. 1A**) and bacterial burdens in the bladders (**Fig. 1B**) were quantified. The Δ*gspD* mutant exhibited decreased binding to the catheter and colonization of the bladder, with approximately 10-fold less colony-forming units (CFU) recovered relative to WT. The catheter binding and bladder burden phenotypes were reverted when *gspD* was complemented, demonstrating that observed phenotypes were due to mutation of the T2SS. These results indicate that the T2SS plays a role in *Acinetobacter* uropathogenesis.

**Figure 1.**
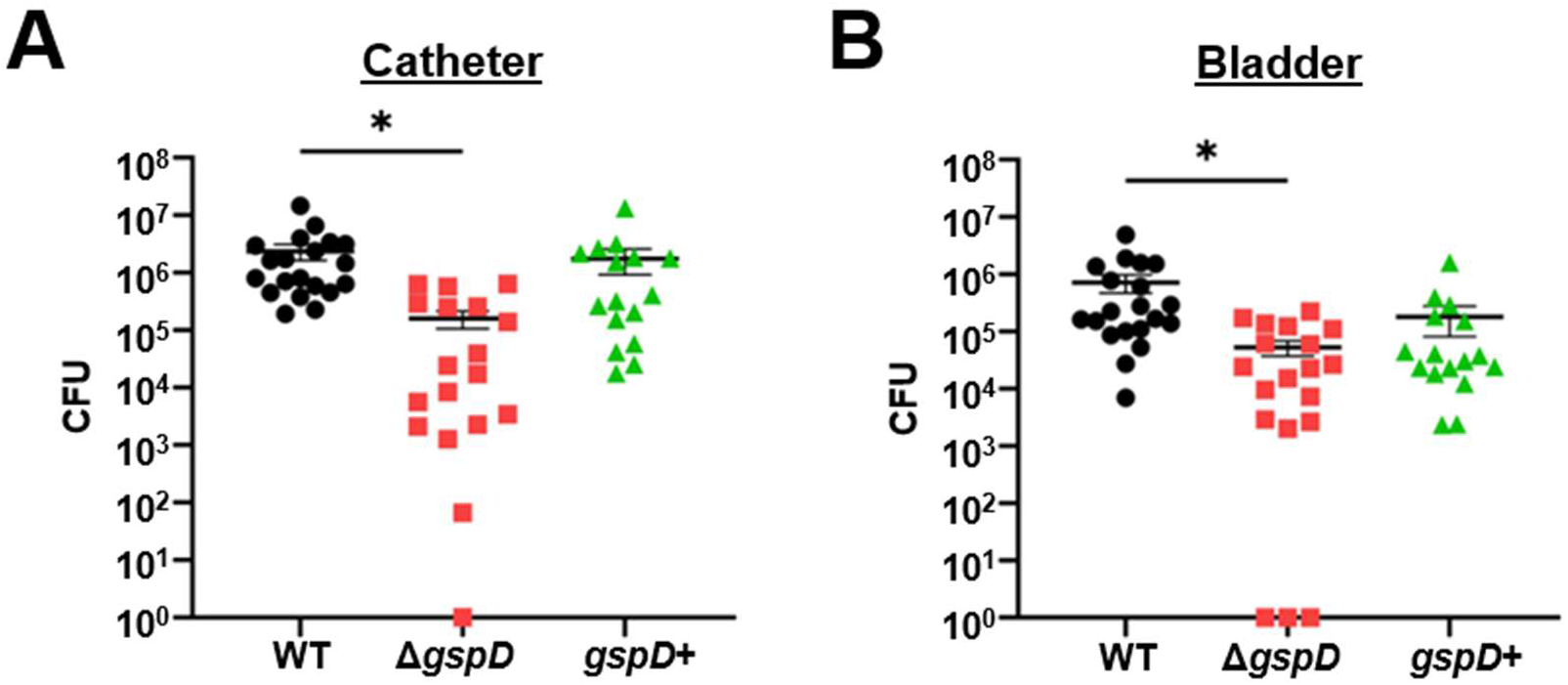
The T2SS is required for full virulence in a murine CAUTI model. Mice were implanted with a catheter followed by transurethral inoculation with UPAB1 WT, Δ*gspD*, or *gspD+* strains. 24 h postinfection, mice were sacrificed, and bacterial burden on the catheter (A) and in the bladder (B) were quantified. Shown are results from at least three pooled experiments. Each data point represents an individual mouse, the horizontal line represents the mean, and the <u>s</u>tandard <u>e</u>rror of the <u>m</u>ean (SEM) is indicated by error bars. \**p*<0.05; One-way ANOVA, Dunnett’s test for multiple comparisons.

### A proteomic analysis identifies a putative T2SS effector with structural similarity to *Yersinia* invasin

To identify T2SS effectors in UPAB1, we performed a proteomic analysis of the supernatant of WT and Δ*gspD* strains. Proteins with decreased abundance with the mutant strain relative to WT were considered putative T2SS effectors (**Table 1**). As expected, we identified orthologs of proteins found in previous *Acinetobacter* T2SS effector screens, including the most differentially identified protein, a 5’-<u>m</u>e<u>t</u>hylthio<u>a</u>denosine (MTA)/<u>S</u>-<u>a</u>denosyl<u>h</u>omocysteine (SAH) nucleosidase, as well as lipases, a CSLREA domain-containing protein, GGT, and the glycoprotease CpaA (24, 28, 29). However, we additionally identified several proteins not found in previous analyses, several of which were annotated as hypothetical proteins of unknown function.

**Table 1.**
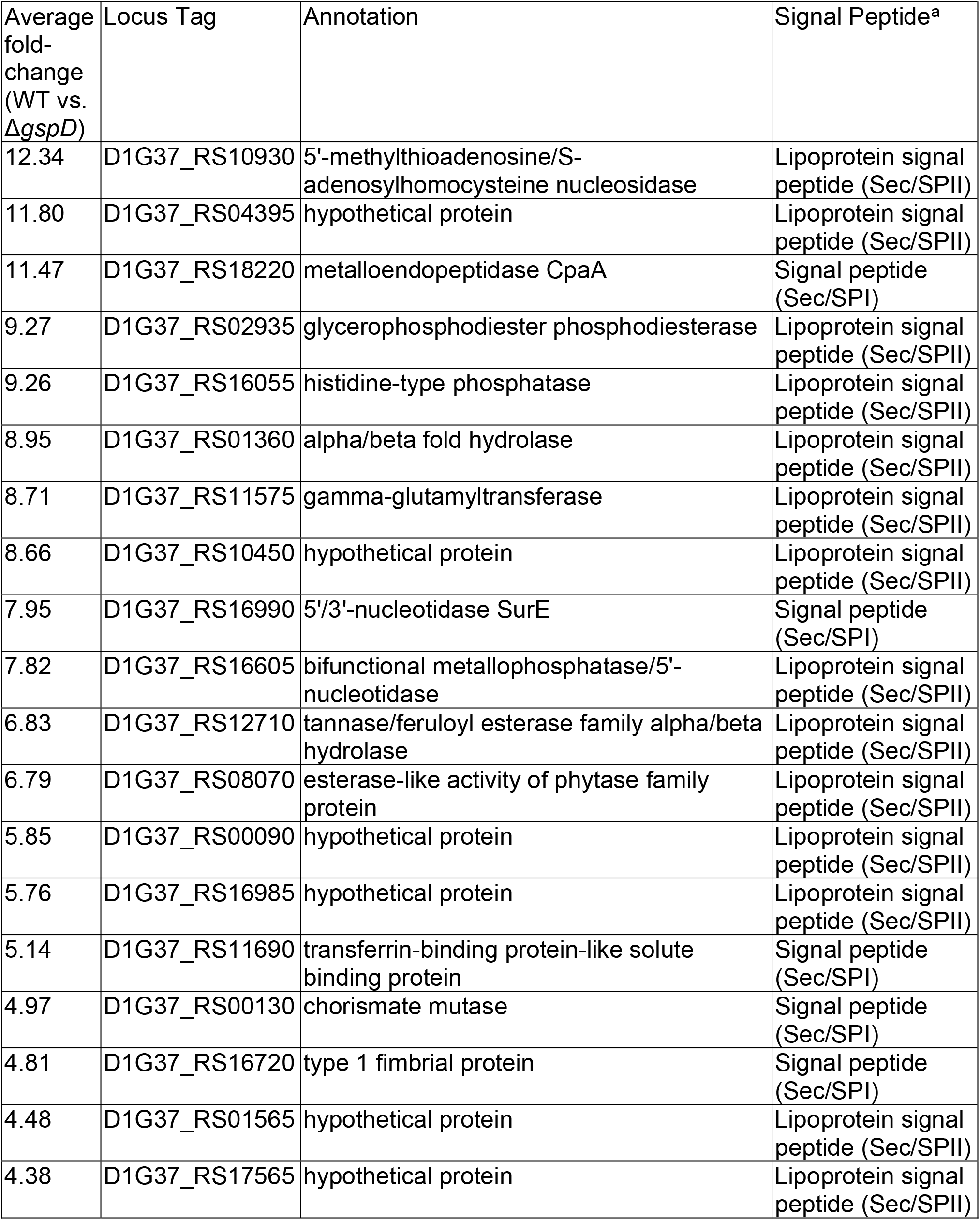
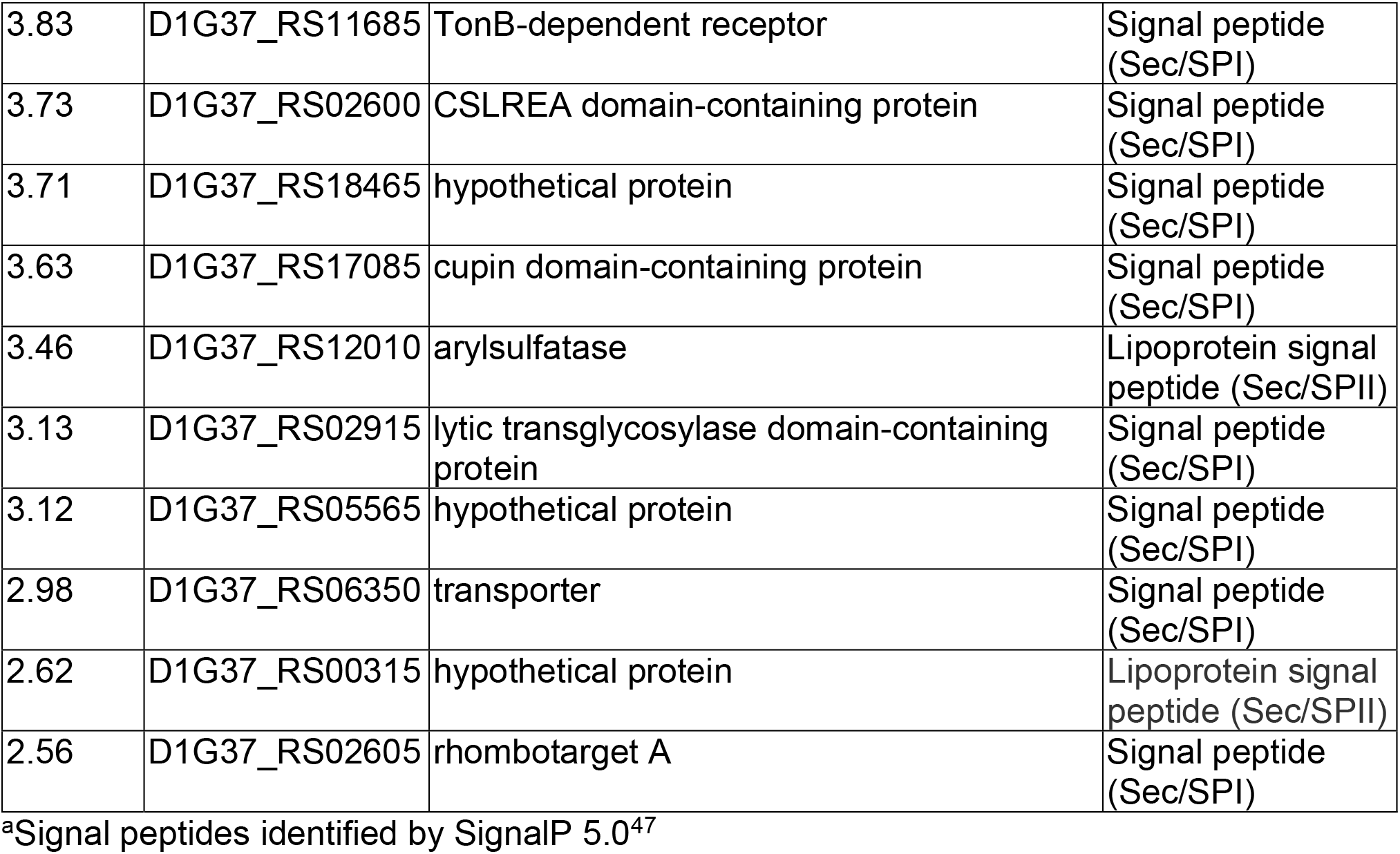
Putative T2SS-dependent secretome of UPAB1

| Average fold-change (WT vs. *ΔgspD*) | Locus Tag | Annotation | Signal Peptide<sup>a</sup> |
| --- | --- | --- | --- |
| 12.34 | D1G37_RS10930 | 5'-methylthioadenosine/S-adenosylhomocysteine nucleosidase | Lipoprotein signal peptide (Sec/SP<sub>II</sub>) |
| 11.80 | D1G37_RS04395 | hypothetical protein | Lipoprotein signal peptide (Sec/SP<sub>II</sub>) |
| 11.47 | D1G37_RS18220 | metalloendopeptidase CpaA | Signal peptide (Sec/SP<sub>I</sub>) |
| 9.27 | D1G37_RS02935 | glycerophosphodiester phosphodiesterase | Lipoprotein signal peptide (Sec/SP<sub>II</sub>) |
| 9.26 | D1G37_RS16055 | histidine-type phosphatase | Lipoprotein signal peptide (Sec/SP<sub>II</sub>) |
| 8.95 | D1G37_RS01360 | alpha/beta fold hydrolase | Lipoprotein signal peptide (Sec/SP<sub>II</sub>) |
| 8.71 | D1G37_RS11575 | gamma-glutamyltransferase | Lipoprotein signal peptide (Sec/SP<sub>II</sub>) |
| 8.66 | D1G37_RS10450 | hypothetical protein | Lipoprotein signal peptide (Sec/SP<sub>II</sub>) |
| 7.95 | D1G37_RS16990 | 5'/3'-nucleotidase SurE | Signal peptide (Sec/SP<sub>I</sub>) |
| 7.82 | D1G37_RS16605 | bifunctional metallophosphatase/5'-nucleotidase | Lipoprotein signal peptide (Sec/SP<sub>II</sub>) |
| 6.83 | D1G37_RS12710 | tannase/feruloyl esterase family alpha/beta hydrolase | Lipoprotein signal peptide (Sec/SP<sub>II</sub>) |
| 6.79 | D1G37_RS08070 | esterase-like activity of phytase family protein | Lipoprotein signal peptide (Sec/SP<sub>II</sub>) |
| 5.85 | D1G37_RS00090 | hypothetical protein | Lipoprotein signal peptide (Sec/SP<sub>II</sub>) |
| 5.76 | D1G37_RS16985 | hypothetical protein | Lipoprotein signal peptide (Sec/SP<sub>II</sub>) |
| 5.14 | D1G37_RS11690 | transferrin-binding protein-like solute binding protein | Signal peptide (Sec/SP<sub>I</sub>) |
| 4.97 | D1G37_RS00130 | chorismate mutase | Signal peptide (Sec/SP<sub>I</sub>) |
| 4.81 | D1G37_RS16720 | type 1 fimbrial protein | Signal peptide (Sec/SP<sub>I</sub>) |
| 4.48 | D1G37_RS01565 | hypothetical protein | Lipoprotein signal peptide (Sec/SP<sub>II</sub>) |
| 4.38 | D1G37_RS17565 | hypothetical protein | Lipoprotein signal peptide (Sec/SP<sub>II</sub>) |
| 3.83 | D1G37_RS11685 | TonB-dependent receptor | Signal peptide (Sec/SPI) |
| 3.73 | D1G37_RS02600 | CSLREA domain-containing protein | Signal peptide (Sec/SPI) |
| 3.71 | D1G37_RS18465 | hypothetical protein | Signal peptide (Sec/SPI) |
| 3.63 | D1G37_RS17085 | cupin domain-containing protein | Signal peptide (Sec/SPI) |
| 3.46 | D1G37_RS12010 | arylsulfatase | Lipoprotein signal peptide (Sec/SPII) |
| 3.13 | D1G37_RS02915 | lytic transglycosylase domain-containing protein | Signal peptide (Sec/SPI) |
| 3.12 | D1G37_RS05565 | hypothetical protein | Signal peptide (Sec/SPI) |
| 2.98 | D1G37_RS06350 | transporter | Signal peptide (Sec/SPI) |
| 2.62 | D1G37_RS00315 | hypothetical protein | Lipoprotein signal peptide (Sec/SPII) |
| 2.56 | D1G37_RS02605 | rhombotarget A | Signal peptide (Sec/SPI) |

<sup>a</sup>Signal peptides identified by SignalP 5.0<sup>47</sup>

The most differentially abundant protein of unknown function in the supernatant of WT and Δ*gspD* strains was D1G37_RS04395. D1G37_RS04395 is a 492 amino acid, 51.92 kDa protein predicted by SignalP 5.0 to contain an N-terminal lipoprotein secretion signal (41). Interestingly, a fold-recognition analysis of D1G37_RS04395 with Phyre^2^ revealed the invasin InvA of *Y. pseudotuberculosis* as the best match (99.7% confidence; 72.0% coverage) (**Fig. S1**) (42, 43). Submission of D1G37_RS04395 to I-TASSER similarly predicted structural similarity to InvA (44–46). Alphafold2 was then used to generate a model of D1G37_RS04395 to compare to the known structure of InvA (**Fig. 2A**) (43, 47). D1G37_RS04395 is predicted to have three IG-like domains and is capped with a C-terminal lectin-like domain that is intimately associated with the most C-terminal IG-like domain. This is comparable to the passenger domain of InvA, with the exception that InvA contains an additional IG-like domain (43). A key difference between D1G37_RS04395 and InvA, however, is that InvA is an autotransporter (T5SS) attached to the membrane by an N-terminal β-barrel domain, whereas D1G37_RS04395 is a putative T2SS effector predicted to be attached to the membrane as a lipoprotein (**Figs. 2B-C**) (35, 36). Given the apparent structural similarity of D1G37_RS04395 to the passenger domain of InvA, we will refer to this protein herein as InvL (<u>inv</u>asin-<u>l</u>ike protein).

**Figure 2.**
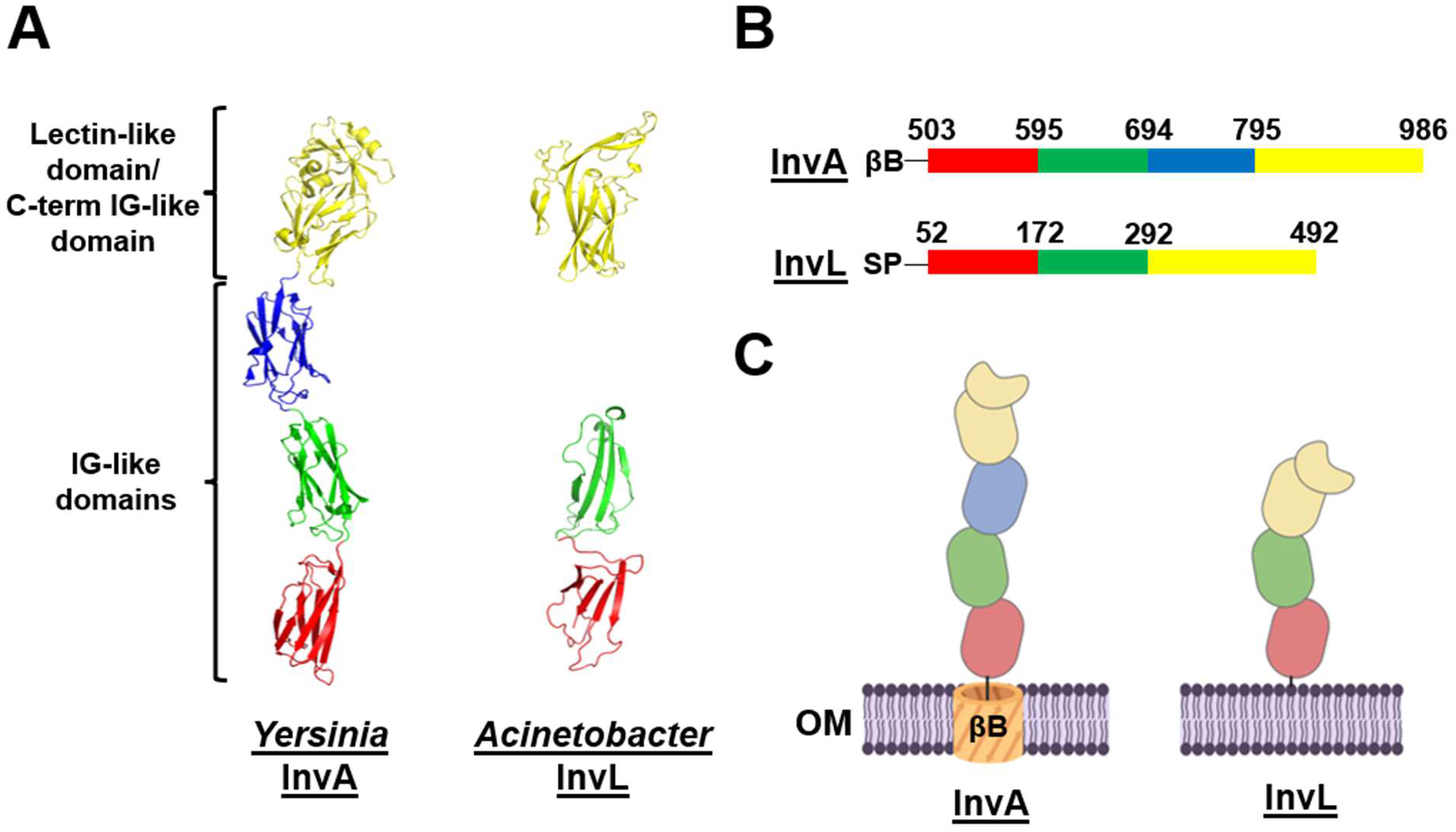
InvL has predicted structural similarity to InvA of *Yersinia*. (A) The crystal structure of the passenger domain of *Yersinia* InvA (PDB: 1CWV) is shown in the left panel, and the AlphaFold2 predicted structure of the analogous region of InvL is pictured in the right panel. (B) Amino acids corresponding to sub-domains of the passenger domain of InvA and the predicted homologous region of InvL. (C) Graphic depiction of the structure of InvA and the predicted structure of InvL; created with BioRender.com. Red, green, and blue colors denote individual IG-like domains, and yellow denotes the C-terminal IG-like domain in intimate contact with the lectin-like domain. βB, <u>β</u>-barrel domain; SP, <u>s</u>ignal <u>p</u>eptide; OM, <u>o</u>uter <u>m</u>embrane.

### The T2SS is required for InvL secretion and surface exposure

To confirm that secretion of InvL is dependent on a functional T2SS, WT, Δ*gspD*, and *gspD+* were transformed with a plasmid expressing *invL* with a His_6_ tag (pBAV-Apr::invL-his_6_). Whole cell lysate and supernatant fractions from the bacteria were then assessed by immunoblot for the presence of InvL-His_6_ (**Fig. 3**). InvL was found in the supernatant of WT and *gspD+* strains, whereas the Δ*gspD* mutant failed to secrete the protein. This confirmed that InvL is a T2SS substrate. Immunoblot analyses demonstrated that, in addition to being released into the supernatant, InvL was cell-associated (**Fig. 3**). As intimin-invasin family proteins function as surface-localized adhesins, we hypothesized that cell-associated InvL is surface-localized. To test this, proteinase K susceptibility assays wherein surface-exposed proteins are degraded and intracellular proteins are left intact were performed with bacteria expressing *invL-his_6_*. In WT UPAB1, InvL-His_6_ was completely degraded with proteinase K treatment, whereas the intracellular control, <u>RNA</u> polymerase (RNAP), was not (**Fig. 4A**). On the contrary, treatment with Triton X-100 and proteinase K resulted in degradation of both InvL-His_6_ and RNAP. These results indicate that InvL is surface-localized, consistent with its putative role as an adhesin. Interestingly, proteinase K failed to degrade InvL-His_6_ expressed in the Δ*gspD* mutant, indicating the protein did not reach the surface of the bacteria (**Fig. 4B**). Alternatively, InvL-His_6_ was readily degraded by proteinase K treatment of *gspD+* (**Fig 4C**), demonstrating that the T2SS is required for InvL surface localization.

**Figure 3.**
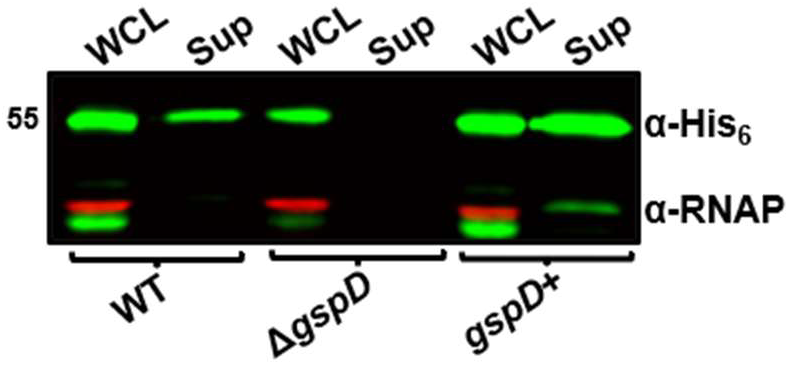
The T2SS is required for InvL secretion. <u>W</u>hole <u>c</u>ell <u>l</u>ysate (WCL) and <u>sup</u>ernatant (Sup) fractions from WT, Δ*gspD*, and *gspD+* UPAB1 cultures harboring *pBAV-Apr::invL-his_6_* were probed for InvL (α-His_6_) and RNAP by immunoblot. Numbers to the left indicate molecular weight in kDa. At least two biological replicates were performed, yielding similar results, and a representative blot from one replicate is shown.

**Figure 4.**
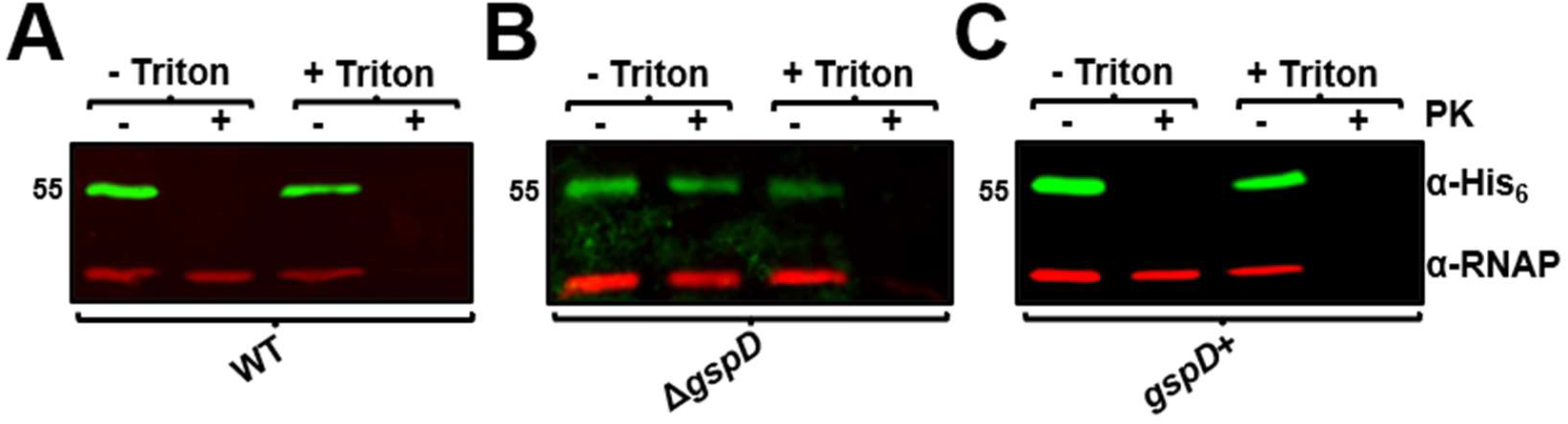
The T2SS is required for InvL surface localization. WT (A), Δ*gspD* (B), and *gspD+* (C) UPAB1 cells harboring *pBAV-Apr::invL-his_6_* were treated with <u>p</u>roteinase <u>K</u> (PK; +) or left untreated (-) in the presence or absence of Triton X-100. Cells were then probed for InvL (α-His_6_) and RNAP by immunoblot. Numbers to the left indicate molecular weight in kDa. At least two biological replicates were performed, yielding similar results, and a representative blot from one replicate is shown.

As InvL is predicted to be a lipoprotein, its identification in the supernatant is somewhat surprising. To further examine the form in which InvL is secreted, insoluble and soluble fractions from the supernatant of WT UPAB1 expressing *invL-his_6_* were separated by ultracentrifugation. Immunoblot analysis of these fractions revealed that InvL was primarily found in the insoluble fraction (**Fig. S2**). This localization could be due to the presence of InvL in either lysis byproducts or possibly <u>o</u>uter <u>m</u>embrane <u>v</u>esicles (OMVs). The physiological relevance of OMVs *in Acinetobacter* pathobiology remains to be investigated and will be the focus of subsequent work.

### InvL binds to extracellular matrix components

We next examined whether InvL could bind to <u>e</u>xtra<u>c</u>ellular <u>m</u>atrix (ECM) components. First, we used <u>e</u>nzyme-<u>l</u>inked <u>i</u>mmunosorbent <u>a</u>ssays (ELISAs) to investigate if InvL binds to α5β1 integrin (**Fig. 5A**) and collagen V (**Fig. 5B**). These molecules are the binding partners of InvA and an *E. coli* homolog, FdeC, respectively (39, 48, 49). We determined that InvL bound to α5β1 integrin and collagen V with dissociation constants (K_dS_) of 0.38 nM and 15 μM, respectively. On the contrary, InvL did not appreciably bind the negative control, <u>b</u>ovine <u>s</u>erum <u>a</u>lbumin (BSA) at the tested concentrations. Since UPAB1 colocalizes with fibrinogen deposited on the catheter during CAUTI, we assessed if InvL interacts with fibrinogen (10). We found that InvL bound to fibrinogen with the highest affinity of all ECM components tested (K_d_=0.19 nM; **Fig. 5C**). InvL, similar to InvA and FdeC, did not exhibit any significant binding to mucin, another common ECM component (**Fig. 5D**).

**Figure 5.**
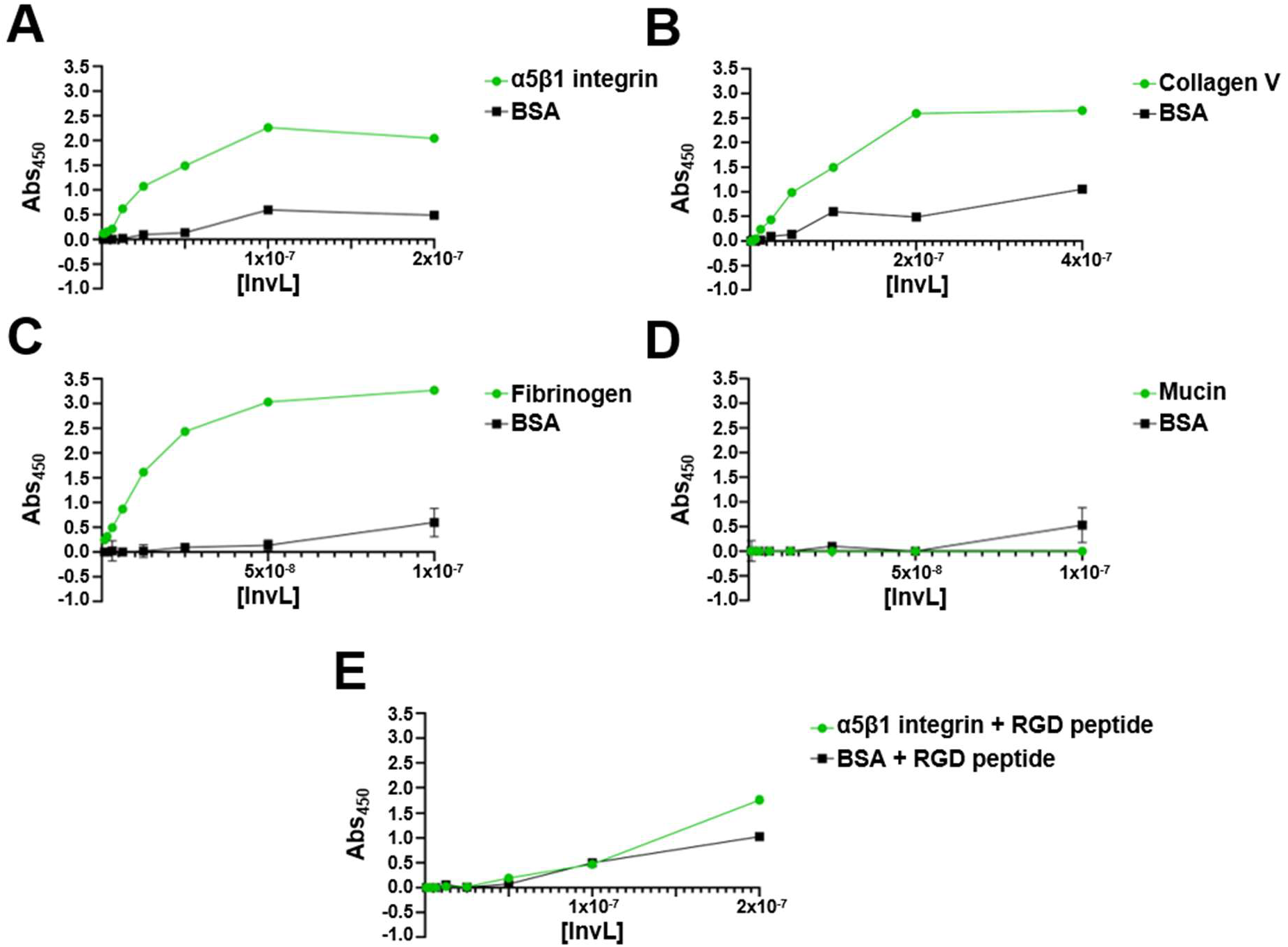
InvL binds to multiple ECM components. ELISAs were performed to assess binding of InvL to α5β1 integrin (A), collagen V (B), fibrinogen (C), mucin (D), or α5β1 integrin in the presence of RGD-containing peptide (E). BSA served as a negative binding control. Shown are the results from two biological replicates, and SEM is indicated by error bars.

The interaction between InvA and β1 integrins is competitively inhibited by arginine-glycine-aspartate (RGD)-containing peptides, a motif required for the interaction of fibronectin with β1 integrin (50–52). RGD-containing peptides also abrogated binding of InvL to α5β1 integrin (**Fig. 5E**), indicating that InvL and InvA likely bind β1 integrins via similar mechanisms. Together, these results demonstrate that InvL binds α5β1 integrin, collagen V, and fibrinogen *in vitro*.

### InvL facilitates UPAB1 adhesion to bladder and kidney epithelial cells

To examine if InvL plays a role in binding to epithelial cells relevant to the CAUTI model, we determined adhesion of UPAB1 WT, Δ*invL*, and *invL+* strains to kidney (MDCK; **Fig. 6A**) and bladder (5637; **Fig. 6B**) cells. The Δ*invL* strain exhibited significantly decreased adhesion to both cell types relative to WT at a <u>m</u>ultiplicity <u>o</u>f infection (MOI) of one, and a similar trend was observed at an MOI of five. This phenotype was reversed in the *invL+* strain, confirming a role for InvL in urinary tract epithelial adhesion.

**Figure 6.**
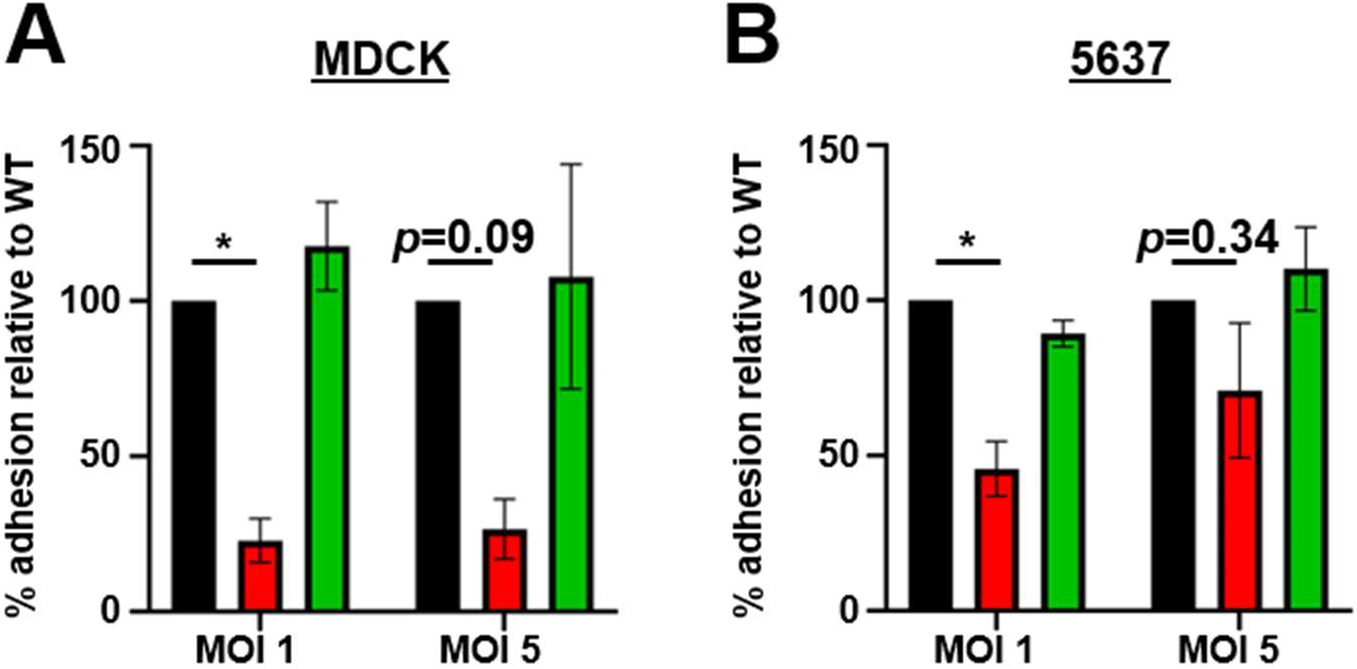
InvL facilitates adhesion to epithelial cells *in vitro*. UPAB1 WT, Δ*invL*, and *invL+* strains were used in adhesion assays with MDCK (A) and 5637 (B) epithelial cells. The mean from three biological replicates is shown and error bars represent the SEM. \**p*<0.05; One-way ANOVA, Dunnett’s test for multiple comparisons.

Because *Yersinia* InvA can mediate epithelial cell invasion, we performed antibiotic protection assays to assess if InvL performs a similar function (53–57). However, internalization was not observed in any of the cell lines tested, and immunofluorescence assays following epithelial cell infection confirmed UPAB1 is not invasive (**Fig. S3**). These results are not surprising given that few publications exist regarding *Acinetobacter* internalization in epithelial cells, and internalization results are highly variable depending on the strain and cell line tested (58–63). Altogether, our results demonstrate a role for InvL in binding to diverse epithelial cell lines, but a possible role in invasion in other strains or cells types cannot be ruled out.

### InvL is required for full virulence in the CAUTI model

Given the role of InvL as an adhesin, we tested if InvL contributes to *Acinetobacter* virulence in the murine CAUTI model. Mice were implanted with a catheter followed by transurethral inoculation with WT, Δ*invL*, or *invL+* strains. 24 h postinfection, mice were sacrificed, and bacteria adhered to the catheters (**Fig. 7A**) and bacterial burdens in the bladders (**Fig. 7B**) were quantified. The Δ*invL* mutant had a significant reduction in catheter binding, exhibiting approximately a 100-fold decrease relative to the WT strain, and this phenotype was reversed by genetic complementation. Δ*invL* also had a significant defect in bladder colonization relative to the WT strain, with more than 10-fold reduced CFU recovered. This phenotype was partially complemented with the *invL+* strain. Overall, these results indicate an important role for InvL in *A. baumannii* uropathogenesis.

**Figure 7.**
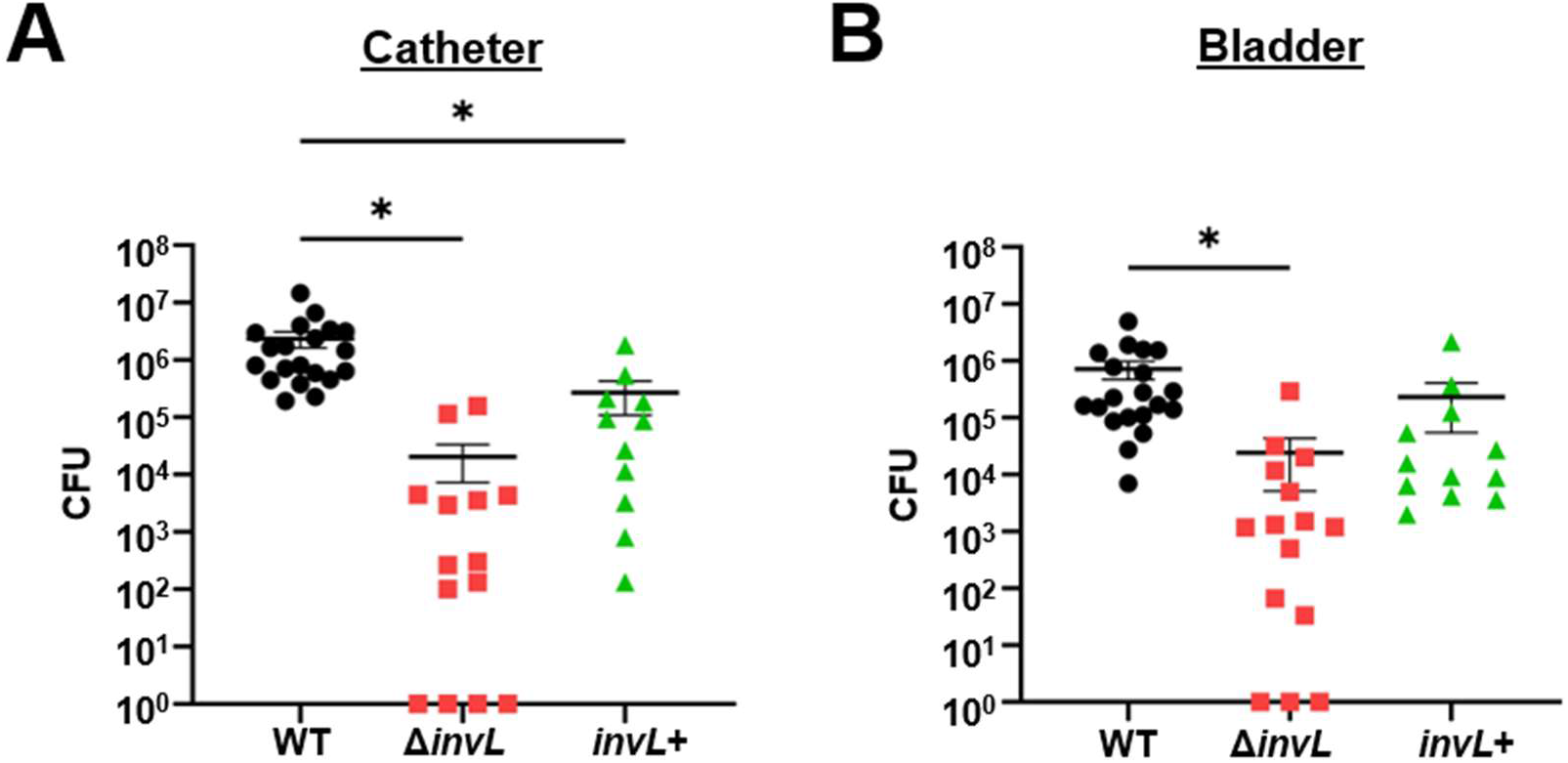
InvL is required for full virulence in a murine CAUTI model. Mice were implanted with a catheter followed by transurethral inoculation with UPAB1 WT, Δ*invL*, or *invL+* strains. 24 h postinfection, mice were sacrificed, and bacterial burden on the catheter (A) and in the bladder (B) were quantified. Shown are results from at least three pooled experiments. Each data point represents an individual mouse, the horizontal line represents the mean, and the SEM is indicated by error bars. \**p*<0.05; One-way ANOVA, Dunnett’s test for multiple comparisons.

### InvL is encoded by ACB complex strains and has high prevalence among international clone 2 isolates

As virulence factors could differ among isolates, we assessed the distribution of InvL across the *Acinetobacter* genus. Specifically, we performed a targeted ortholog search for *invL* and adjacent genes in 3052 available genomes from the <u>N</u>ational <u>C</u>enter for <u>B</u>iotechnology Information (NCBI) RefSeq Database (64). In genomes where InvL is encoded, the corresponding gene is flanked by genes encoding a TIGR01244 family phosphatase (D1G37_04390 in UPAB1) and a dihydrolipoyl dehydrogenase (D1G37_RS04400 in UPAB1) (**Fig. 8A**). While the microsynteny of the two flanking genes is conserved throughout ACB complex, only 74% (2029/2728) of the isolates additionally harbor the *invL* gene. In 23% (641/2728), a second cluster variant exists, where the *invL* gene is replaced by a gene encoding a hypothetical protein (A1S_3863 in ATCC 17978). The remainder of isolates (3%; 58/2728) have no/another intervening gene. The gene represented in the second cluster variant is located on the opposite strand and shares no significant sequence similarity with InvL. Outside the ACB complex, the gene cluster is represented in only 216/324 (67%) of the isolates. Of these, again the majority encode *invL* (157/216; 72%). The second cluster variant is present in 32/216 (15%) of the isolates, and the remaining 13% have either no or another intervening gene. Of note, InvL distribution correlates with international clone (IC) number, as nearly all strains belonging to IC-2, IC-4, and IC-5 groups encode InvL (**Fig. 8B**). Alternatively, strains belonging IC-1, IC-6, and IC-8 predominantly encode the second cluster variant. As IC-2 clones are among the most commonly isolated in clinical settings, the prevalence of InvL in these strains is intriguing (1–3, 65, 66). However, the fact that other international clones which are associated with human infection do not predominantly encode InvL (e.g., IC-1) could imply that some other protein(s) may compensate for the absence of this adhesin. In sum, we conclude that InvL is prevalent in the ACB complex and is found in nearly all IC-2 strains, which are relevant to human disease.

**Figure 8.**
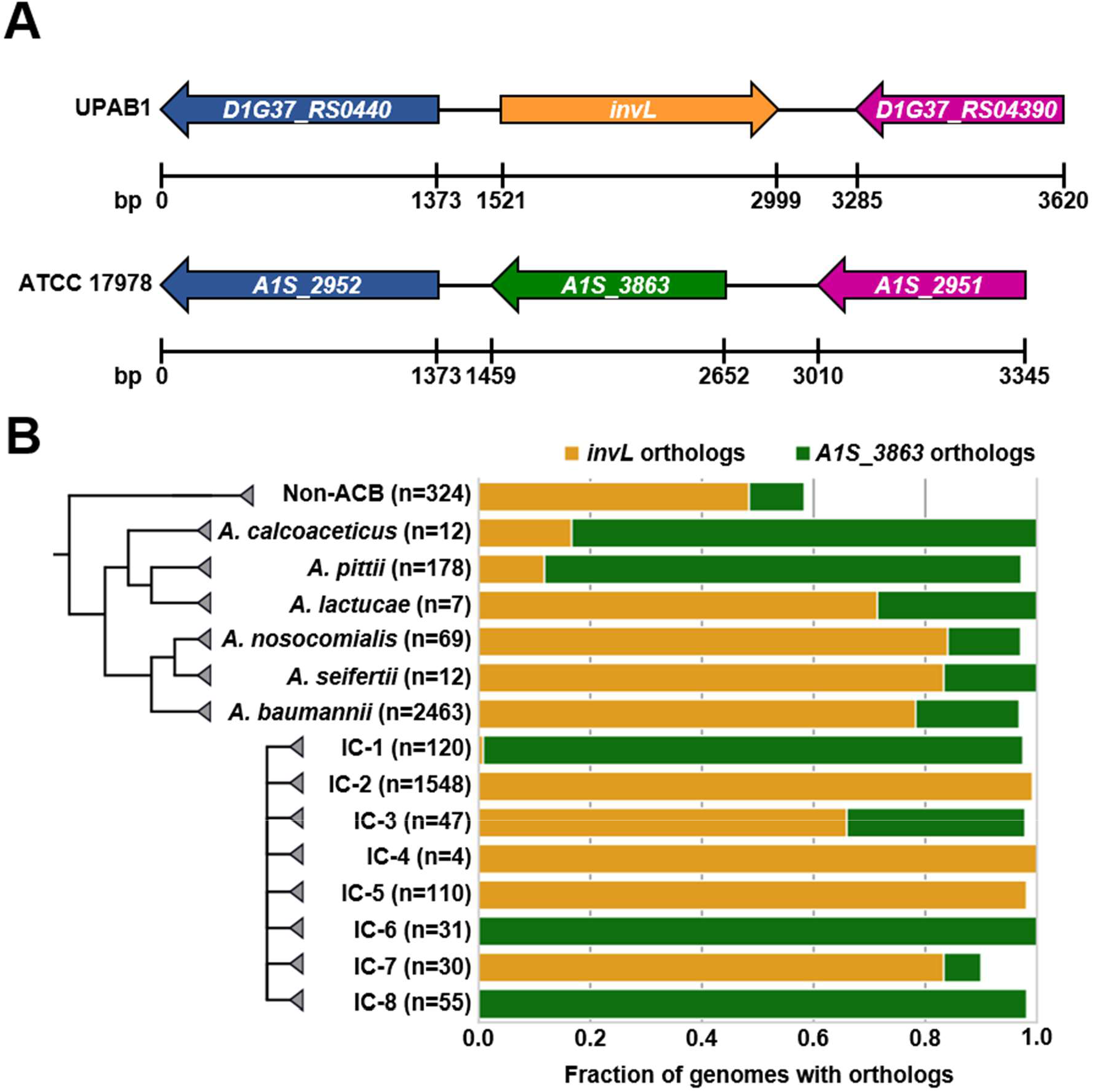
InvL is encoded by ACB complex strains and has high prevalence in IC-2 clones. (A) InvL is encoded at a common locus in ACB complex strains. Orthologs of of InvL (top) and orthologs of A1S_3863 (bottom) are encoded at a common locus between genes encoding a dihydrolipoyl dehydrogenase (D1G37_RS0440 in UPAB1 and A1S_2952 in ATCC 17978; blue) and a TIGR01244 family phosphatase (D1G37_RS04390 in UPAB1 and A1S_2951 in ATCC 17978; purple). (B) Distribution of *invL* and *A1S_3863* in *Acinetobacter*. The fraction of genomes encoding InvL and A1S_3863 are reported by ACB species and international clone number.

## DISCUSSION

*Acinetobacter* has emerged as a pathogen of significant clinical importance due to the increasing frequency of multidrug resistant strains identified (6, 8, 9). Approximately 20% of isolates are derived from the urinary tract, highlighting the uropathogenic potential of *A. baumannii* (10, 11). Mutation of the T2SS in the uropathogenic strain UPAB1 resulted in decreased catheter-binding and lower burden in the bladder, indicative of a key role during urinary tract virulence. Through a proteomic analysis, we identified a surface-exposed T2SS effector that belongs to the intimin-invasin family, referred to here as InvL. We found that recombinant InvL binds to host ECM components and that the UPAB1 Δ*invL* mutant strain has a significant defect in binding to bladder and kidney epithelial cells. Accordingly, the Δ*invL* mutant was attenuated in the CAUTI model. In all, our study demonstrates the importance of the T2SS for *Acinetobacter* uropathogenesis and describes a novel T2SS effector, InvL, as a previously unrecognized adhesin and virulence factor that is encoded by most clinical isolates.

Interestingly, although protein modeling revealed structural similarity of InvL to *Yersinia* InvA, InvL is unique compared to other invasin homologs. First, while canonical invasin homologs are T5SSs, outer membrane localization of InvL is dependent on the T2SS. To our knowledge, this is the first report of an intimin-invasin family T2SS effector. Second, unlike other InvA homologs which are anchored to the outer membrane via a large N-terminal β-barrel domain, InvL is predicted to be a lipoprotein based on the presence of a lipoprotein secretion signal (36, 41). While the T2SS is known to secrete soluble effectors into the extracellular milieu, it has also been reported to be essential for surface localization of some lipoproteins similar to what is reported here for InvL (67). One of the best characterized examples is the T2SS-dependent surface localization of the lipoprotein pullulanase (PulA), a starch-debranching enzyme of *Klebsiella oxytoca* (68–70). PulA, like many other lipoproteins, is translocated from the cytoplasm to the periplasm via the sec pathway, where processing and acylation occurs to attach the protein to the periplasmic leaflet of the inner membrane (69, 71). Then, whereas most outer membrane lipoproteins of Gram-negative bacteria are transported from the inner membrane to the outer membrane via the localization <u>o</u>f <u>l</u>ipoproteins (Lol) pathway, PulA uses the T2SS machinery (68–70). The mechanism by which the T2SS is able to recognize its lipoprotein effectors and transport lipidated proteins across the periplasm and outer membrane remains elusive. However, this report of InvL adds to the growing list of known T2SS effectors which are surface exposed lipoproteins (67). As several of these proteins are involved in pathogenesis and key metabolic functions (e.g., SslE of *E. coli* and MtrC and OmcA *Shewanella oneidensis)*, a better understanding of the mechanism of T2SS-dependent lipoprotein transport is needed (67, 72–77).

Similar to other members of the intimin-invasin protein family, InvL is able to directly bind to ECM components and facilitate cell adhesion. Specifically, InvL binds with high affinity to α5β1 integrin, collagen V, and fibrinogen. α5β1 integrin and collagen V are also bound by the intimin-invasin family proteins InvA and FdeC of *Yersinia* and *E. coli*, respectively (39, 49). Interestingly though, InvA and FdeC appear to bind to a single defined ECM component, indicating that InvL has a broader specificity. While domains/residues required for the FdeC-collagen interaction have not been defined, the interaction between InvA and β1 integrins has been extensively studied (48, 78–82). Binding of InvA to β1 integrins specifically requires the C-terminal lectin-like domain, which is present in InvL and absent in FdeC (49, 80, 82). It is tempting to speculate that the predicted InvL lectin-like domain could be involved in binding to α5β1 integrin and that characteristics of Ig-like domains may be responsible for recognition of collagen V. However, it should be noted that residues known to be important for the interaction between InvA and β1 integrins do not appear to be conserved in InvL (78–81). This is despite the observation that the interactions of InvL and InvA with α5β1 integrin can be similarly inhibited by RGD peptide (50–52). The low sequence identity between InvL and invasin homologs makes it difficult to accurately predict key regions of the protein required for recognition of ECM components. The structural characterization of InvL will, in the future, provide mechanistic insights into the interaction between InvL and the ECM.

Our characterization of InvL adds to the growing list of known *Acinetobacter* adhesins that facilitate binding to epithelial cells, which includes Type IV pili, the Ata autotransporter, the FhaB/FhaC two-partner secretion system, and <u>b</u>iofilm-<u>a</u>ssociated <u>p</u>rotein (Bap) (83–93). ECM components bound by Type IV pili and Bap are not known. However, Ata has been shown to bind with high affinity to collagen (types I, III, IV, and V), laminin, and glycosylated fibronectin, and FhaBC reportedly binds to fibronectin (85, 88, 89, 92). While the collagen-binding role of InvL may be redundant with that of Ata, InvL is the first published *Acinetobacter* adhesin to bind directly to α5β1 integrin or fibrinogen. Additionally, InvL is now the first known adhesin to be important for *A. baumannii* UTIs. Future work will determine the roles of these known adhesins in the multiple niches colonized by this bacterium.

While we have established that InvL is a virulence factor involved in *Acinetobacter* uropathogenesis, the mechanism(s) by which InvL participates in CAUTI is unknown. The fact that we identified fibrinogen as the highest-affinity binding partner for InvL is particularly intriguing with respect to CAUTI. Tissue damage from catheterization results in an inflammatory response that leads to increased levels of fibrinogen in the bladder which ultimately coats the catheter (94, 95). We recently showed that *Acinetobacter*, similar to other uropathogens such as *Enterococcus faecalis* and *Staphylococcus aureus*, colocalizes with fibrinogen on the catheter (10, 96, 97). This interaction is dependent on the virulence factors EbpA and ClfB in *E. faecalis* and S. *aureus*, respectively (95, 96). It is tempting to speculate that InvL may serve a similar function with respect to *Acinetobacter* catheter binding. However, future work is required to examine if InvL mediates catheter-binding via fibrinogen or if the ability of InvL to associate with other ECM components is responsible for its role in the CAUTI model. Additionally, future work is required to determine if InvL similarly serves as a virulence factor in other infection models (e.g., pneumonia and bacteremia). Ultimately, a better understanding of *A. baumannii* virulence factors such as InvL may aid in the development of novel therapeutics to combat infection by these increasingly multidrug resistant bacteria.

## MATERIALS AND METHODS

### Bacterial plasmids, strains, and growth conditions

Plasmids and strains used in this study are detailed in **Table S1**. Cultures were grown at 37°C using Lennox broth/agar unless otherwise noted. The following antibiotic concentrations were used when appropriate; 10 μg/ml chloramphenicol, 100 μg/ml ampicillin, 30 μg/ml apramycin, or 50 μg/ml zeocin for *E. coli* and 10 μg/ml chloramphenicol, 50 μg/ml apramycin, 50 μg/ml zeocin, or 300 μg/ml hygromycin B for *A. baumannii*.

### Generation of constructs used in this study

Plasmids and primers used in this study are detailed in **Table S1** and **Table S2**, respectively. Detailed information for construction of plasmids and strains used in the study is provided in the supplementary materials.

### Murine infection experiments

Animal experiments were approved by the Washington University Institutional Animal Care and Use Committee. The murine CAUTI model was performed as previously described (10). Briefly, 6- to 8-wk-old female C57BL/6 mice (Charles River Laboratories, Wilmington, MA) anesthetized with 4% isoflurane were implanted transurethrally with a 4- to 5-mm piece of silicon tubing (catheter). Once-passaged static grown bacterial strains were washed twice and resuspended in phosphate-buffered saline (PBS). Mice were then transurethrally inoculated with 50 μl of the bacterial suspension containing 2 x 10^8^ CFU. 24 h postinfection, mice were sacrificed, and bacterial load on catheters and in bladders were quantified by serial dilution plating.

### Proteomic analysis of the T2SS-dependent secretome

Bacterial culture conditions, secreted protein enrichment, digestion of secretome samples, and liquid chromatography-mass spectrometry analysis of secretome samples was performed as previously described (98). Details are provided in the supplementary materials. The resulting MS data and search results have been deposited into the PRIDE ProteomeXchange Consortium repository and can be accessed using with the accession numbers PXD030460 (Username; Password: WuMcxzwD) and PXD030491 (Username; Password: 3q2QpAlz) (99, 100).

### Immunoblot analyses

Bacterial whole cell lysate preparation and TCA precipitation of supernatant fractions from mid-exponential cultures was performed as previously described (24, 101). Indicated fractions were separated by SDS-PAGE, transferred to a nitrocellulose membrane, and proteins of interest were probed using polyclonal rabbit anti-His_6_ (1:2000; Invitrogen, Waltham, MA) and monoclonal mouse anti-RNAP (1:3500; BioLegend, San Diego, CA). IRDye-conjugated anti-mouse IgG and anti-rabbit IgG were used as secondary antibodies (1:5000 for each; LI-COR Biosciences, Lincoln, NE), and blots were visualized with the Odyssey CLx imaging system (LI-COR Biosciences).

### Proteinase K accessibility assays

Proteinase K accessibility assays were performed as previously described with some modification (102). Stationary phase cultures were pelleted and resuspended in PBS supplemented with 5 mM MgCl2 to 2.5 OD600. Bacterial suspensions were treated with 200 μg/ml proteinase K (GoldBio, Olivette, MO) or sham treated with an equal volume of diH_2_O in the presence or absence of 2% Triton X-100 (Sigma-Aldrich, St. Louis, MO) at 37°C for 15 minutes. Digestion was terminated by addition of 1 mM phenylmethylsulfonyl fluoride (Amresco LLC, Salon, OH), Laemmli buffer was added to a concentration of 1X, and samples were boiled for 10 minutes. 10 μl of these preparations were separated by SDS-PAGE, and immunoblotting was performed as described above.

### Expression and purification of recombinant InvL

To generate recombinant protein for antibody generation, full-length InvL with a C-terminal His10 tag was expressed from *pET-22b(+)::invL-his10* in Rosetta-gami 2(DE3) cells (Novagen, Madison, WI) using ZYM-5052 autoinducible media for 72 h at 20°C (103). Cells were then lysed using a CF1 cell disrupter (Constant Systems Ltd, Daventry, UK). Following cell lysis, recombinant InvL was purified from the insoluble fraction by Ni-NTA affinity chromatography as previously described (104). Purified protein was then used by Antibody Research Corporation (St. Peters, MO) to generate polyclonal rabbit antibody.

To generate soluble recombinant InvL for ELISAs (see below), InvL lacking the N-terminal signal sequence (as determined by SignalP 5.0) was expressed with a C-terminal His10 tag from pET-22b(+)::invL-his10(-SS) in Rosetta-gami 2(DE3) cells by addition of 1 mM isopropyl-β-d-thiogalactopyranoside for 3 hrs at 37°C (41). Cells were then lysed as above, and recombinant InvL was purified from the soluble fraction using the same procedure as previously described with the exception that buffers were not supplemented with 0.3% N-lauryl-sarcosine (104).

### ELISAs

ELISAs were based on a previously described protocol (49). Detailed protocol information can be found in the supplemental materials.

### Eukaryotic cell adhesion assays

Cell adhesion assays were performed with MDCK and 5637 cells cultured in <u>D</u>ulbecco’s <u>M</u>odified <u>E</u>agle <u>M</u>edium (DMEM) and <u>R</u>oswell <u>P</u>ark <u>M</u>emorial <u>I</u>nstitute <u>1640</u> (RPMI-1640) (Gibco, Dublin, Ireland) supplemented with 10% heat-inactivated fetal bovine serum, respectively. 10^5^ MDCK and 3 x 10^5^ 5637 cells (amounts determined to form confluent monolayers) were plated in 48-well Nunclon Delta cell culture–treated surface plates (ThermoFisher Scientific) overnight at 37°C with 5% CO_2_. Stationary *A. baumannii* strains were pelleted at 5432 × *g* for 5 min and resuspended in PBS. Following one more spin, bacteria were resuspended in the appropriate cell culture media, and 500 μl of suspension at the indicated MOI was added to the appropriate wells. Plates were then centrifuged at 200 × *g* and incubated at 37°C with 5% CO2 for 1 h to allow adhesion. Wells were subsequently washed three times with 500 μl PBS, cells were resuspended in 0.05% Triton X-100 in PBS, and serial dilution plating was performed to quantify bacteria. Concurrently, to quantify intracellular bacteria after the 1 h incubation and washing, the appropriate cell culture medium supplemented with 50 μg/ml colistin was added for 1 h. After washing in PBS, cells were resuspended in 0.05% Triton X-100 in PBS, and serial dilution plating was performed to quantify bacteria.

### Bioinformatic analysis of InvL distribution

Ortholog searches were performed across a database of 3052 Acinetobacter genomes obtained from NCBI RefSeq as previously described (64). Briefly, the collection of orthologous groups that resulted from an all vs all ortholog search using OMA were screened to associate the group that harbors a particular protein sequence of interest (105). The sequences of the associated group collectively served to train a model for the targeted ortholog search across the full database of genomes. The proteins of interest comprise the translated sequences of D1G37_RS04395 (InvL), D1G37_RS0440, D1G37_RS04390, and A1 S_3863. Since the subset of genomes used by Djahanschiri et al. did not include the strain UPAB1, the orthologs identified in the strain MDR-TJ via a reciprocal best blast hit approach were used to associate orthologous groups (64). The resulting presence/absence matrix together with the genus’ phylogeny served as the input for Vicinator v0.32 (https://github.com/BIONF/Vicinator) to trace the microsynteny of the InvL locus based on MDR-TJ and ATCC 17978 as the reference genomes.

## Data availability

Mass spectrometry data has been deposited into PRIDE ProteomeXchange Consortium repository as described above, and all other data is provided within the text or supplementary materials.

## Statistical methods

All statistical analyses were performed using GraphPad Prism version 9.

## ACKNOWLEDGEMENTS

This work was supported by funding to MFF (R01AI144120) and CJL (T32AI007172) through the National Institute of Allergy and Infectious Diseases of the National Institutes of Health. The content is solely the responsibility of the authors and does not necessarily represent the official views of the National Institutes of Health. This study was additionally supported by a grant by the German Research Foundation (DFG) in the scope of the Research Group FOR2251 “Adaptation and persistence of A. baumannii.” (Grant ID EB-285-2/2) to IE. We acknowledge the Molecular Microbiology Imaging Facility at Washington University in St. Louis and Wandy Beatty for confocal microscopy assistance. We also thank Siyuan Ding and Carolina Lopez for providing eukaryotic cell lines used in this manuscript.

